# Investigating endothelial cell transduction and hexon:PF4 binding of ChAdOx1 in the context of VITT

**DOI:** 10.1101/2025.03.09.642192

**Authors:** Charlotte Lovatt, Lars Frängsmyr, Emma A. Swift, Rosie M. Mundy, Alan L. Parker

**Author notes:** **<u>Correspondence:</u> Professor Alan Parker**, Division of Cancer and Genetics, Cardiff University, Heath Park, Cardiff CF14 4XN.

## Abstract

**Background:** Vaccines against SARS-CoV2 have been essential in controlling COVID-19 related mortality and have saved millions of lives. Adenoviral (Ad) based vaccines have played an integral part in this vaccine campaign, with licensed vaccines based on the simian Y25 isolate (Vaxzevria, Astrazeneca) and human Ad type 26 (Jcovden, Janssen) widely adopted. As part of the largest global vaccination programme ever undertaken, ultrarare thromboembolic events have been described in approximately 1:200,000 vaccinees administered with Ad based SARS-CoV2 vaccines.

**Objectives:** The mechanism underpinning these adverse events remain to be completely delineated, but is characterised by elevated autoantibodies against PF4 which, in complex with PF4, cluster, bind FCγRIIa on platelets and induce thrombus formation. Here we investigated the ability of ChAdOx1 to transduce and activate endothelial cells.

**Methods:** Using protein sequence alignment tools and *in vitro* transduction assays, the ability of ChAdOx1 to infect endothelial cells was assessed. Furthermore, the ability of ChAdOx1 infection to activate endothelial cells was determined. Finally, using surface plasmon resonance we assessed the electrostatic interactions between the ChAdOx1 hexon and PF4.

**Results and Conclusions:** Despite lacking the primary cell entry receptor, Coxsackie and Adenovirus Receptor (CAR), ChAdOx1 efficiently transduced endothelial cells in a CAR-independent manner. This transduction did not result in endothelial cell activation. Purified hexon protein from ChAdOx1 preps did, however, bind PF4 with a similar affinity to that previously reported for the whole ChAdOx1 capsid. These data confirm the need to develop non-PF4 binding adenoviral capsids to reduce the potential adverse events associated with VITT.

## Introduction

The ChAdOx1 nCoV-19 vaccine (Vaxzevria/AstraZeneca), derived from chimpanzee adenovirus Y25[1] is credited with saving millions of lives during the COVID pandemic. However, vaccine-induced immune thrombotic thrombocytopenia (VITT) was identified as an ultrarare but serious adverse effect of adenoviral vector-based COVID-19 vaccines[2]. VITT resembles heparin-induced thrombocytopenia (HIT), an autoimmune indication caused by the formation of antibodies against platelet factor 4 (PF4). Patients present with severe thrombocytopenia and thrombosis at uncommon sites, including the cerebral venous sinus, 5-24 days post-vaccination[3]. Existing evidence suggests PF4 forms an electrostatic interaction with the negatively charged ChAdOx1 viral capsid within the inter-hexon space, inducing autoimmune anti-PF4 responses and production of anti-PF4 antibodies[4]. These antibodies form PF4 immune complexes which bind Fc receptors on platelets and neutrophils, resulting in activation and thrombus formation[5].

A contributing mechanism for VITT may involve the inappropriate transduction and activation of endothelial cells (EC) by ChAdOx1. Investigations of ChAdOx1 vector-host interactions are critical to improving understanding of how these rare adverse events occur; therefore, we assessed the direct effects of ChAdOx1 infection of EC. We demonstrate that ChAdOx1 transduces EC despite the absence of their primary entry receptor coxsackie and adenovirus receptor (CAR), but that this does not induce EC activation, suggesting that direct infection of EC by ChAdOx1 does not contribute to the pathogenesis of VITT. Using surface plasmon resonance (SPR) we confirm PF4 binding to purified ChAdOx1 hexon protein, supporting findings that VITT pathogenesis results from electrostatic interactions between PF4 and ChAdOx1.

## Materials and Methods

### Cell lines

Chinese hamster ovary (CHO)-CAR and CHO-BC1 cells expressing CAR and CD46, respectively and human vascular EC were maintained in DMEM-F12 medium (Gibco) supplemented with 10% foetal bovine serum, 1% L-Glutamine and 2% penicillin and streptomycin.

### Viruses

ChAdOx1-GFP, ChAdOx1-S, HAdV-C5, HAdV-C5-26K and HAdV-C5-26K were propagated and purified using the CsCl gradient method [4].

### Cell receptor staining

Cells were stained for receptors using primary CAR RmcB (Millipore, 05-644; 1:500), CD46 (GeneTex, MEM-258; 1:100) and CD31 (Biolegend, 303101; 1:100) antibodies and Alexa-Fluor647-conjugated secondary antibody (ThermoFisher, A21237; 1:1000), then fixed in 4% paraformaldehyde with staining detected using a BD Accuri C6 cytometer. Analysis was performed in FlowJo.

### Fiber knob protein alignment

Sequences for HAdV fiber knob proteins were obtained from NCBI and aligned using Clustal Omega [6].

### Generation of recombinant fiber knob proteins

Recombinant fiber knob proteins were produced as previously described [4].

### Competitive inhibition assays with recombinant fiber knob proteins

Competitive inhibition assays with recombinant fiber knob proteins were performed as previously described[4].

### Viral transduction assay

2.5×10^4^ cells/ well were transduced with viruses in serum-free medium for 3h. Virus was removed and replaced with complete medium for 45h. For GFP transduction, cells were washed, fixed and analysed as described[4]. Luciferase transduction was analysed using Luciferase Assay System kit (Promega), normalised to protein. Luciferase was quantified using a BioTek Cytation 4 plate reader.

### Neuraminidase assay

Neuraminidase transduction assays were performed as above, with seeded cells treated with 50 mU/mL neuraminidase for 1h at 37°C before transduction for 1h on ice.

### Thrombosis assay

EC were transduced as above. Supernatants were collected 6h, 12h, 24h and 48h post-infection, centrifuged to remove debris and EC production of factor IX, interleukin-6 (IL-6), IL-8, plasminogen activator inhibitor-1 (PAI-1), P-selectin, P-selectin glycoprotein ligand (PSGL), sCD40L and tissue factor (TF) was quantified using the LEGENDplex human thrombosis panel (Biolegend) per manufacturer’s instructions.

### Hexon purification

Adenovirus hexons were propagated and purified using the CsCl gradient method [4] with the modification of extracting the top band, containing the empty capsid, rather than the lower band containing complete virions. Anion exchange was performed using an ÄKTA FPLC (Cytvia) using 5 ml HiTrap Q FF columns (Cytvia). Using a 5 ml loop, the extracted band was loaded onto the column and 4 column volumes (CV) of Buffer A (20 mM Hepes; pH 7.4) were run across the column. Hexon was eluted using a linear gradient over ∼10 CV of Buffer B (20 mM Hepes; pH 7.4; 1 M NaCl). Fractions were collected and those corresponding to hexon sized peaks were run on a gel and stained with Coomassie blue (ThermoFisher) to identify the fractions containing hexon protein. Fractions containing hexon were then concentrated with a 50 kDa MWCO (Amicon) centrifugal filter to a final volume of 1 ml and size exclusion chromatography (SEC) performed with a Superose increase 10/300 GL SEC column (Cytvia) on an ÄKTA FPLC (Cytvia). The column was equilibrated with GF buffer (20 mM Hepes; pH 7.4; 150 mM NaCl) and the concentrated fractions were loaded using a 1ml loop. Buffer was then run at 0.5 ml/min and samples collected from 10 ml to 25 ml, corresponding to the hexon peak.

### Surface plasmon resonance

All SPR experiments were performed at 25°C in 10 mM phosphate buffer, pH 7.4, 140 mM NaCl, and 0.27 mM KCl running buffer. Data was collected with a Biacore T-200 instrument at a rate of 1 Hz. Hexons were coupled to the CM5 sensor chip by amine coupling reactions for an immobilization density of ∼1000 resonance units (RU). PF4 was serially diluted in running buffer (0.125 mM, 0.25 mM, 0.5 mM, 1 mM and 2mM) and then injected over the surfaces. Blank samples contained only running buffer. After each cycle, the biosensor surface was regenerated with a 60s pulse of 10 mM Tris-glycine (pH 1.5) at a flow rate of 30 μL/min.

### Statistics

Results are presented as mean ± standard deviation; IC_50_ curves were calculated and fitted by non-linear regression. All statistics and graphs were generated using GraphPad Prism.

## Results and Discussion

### Determining the mechanism by which ChAdOx1 infects CAR-negative EC

The primary virus-cell interaction during infection is mediated by the adenovirus fiber knob, where CAR is a high affinity ChAdOx1 receptor[4]. Alignment of ChAdOx1 fiber knob with other CAR-binding adenoviruses[7],[8],[9] identified shared contacts (Figure 1ai). The use of CAR as a high affinity ChAdOx1 fiber knob receptor was confirmed with competitive inhibition assays with HAdV-C5 and ChAdOx1 demonstrating IC_50_ =0.0457 and 0.0757μg/10^5^ cells respectively (Figure 1_ii_). Cross-species utilisation of CAR has been reported previously for canine adenoviruses[10].

**Figure 1.**
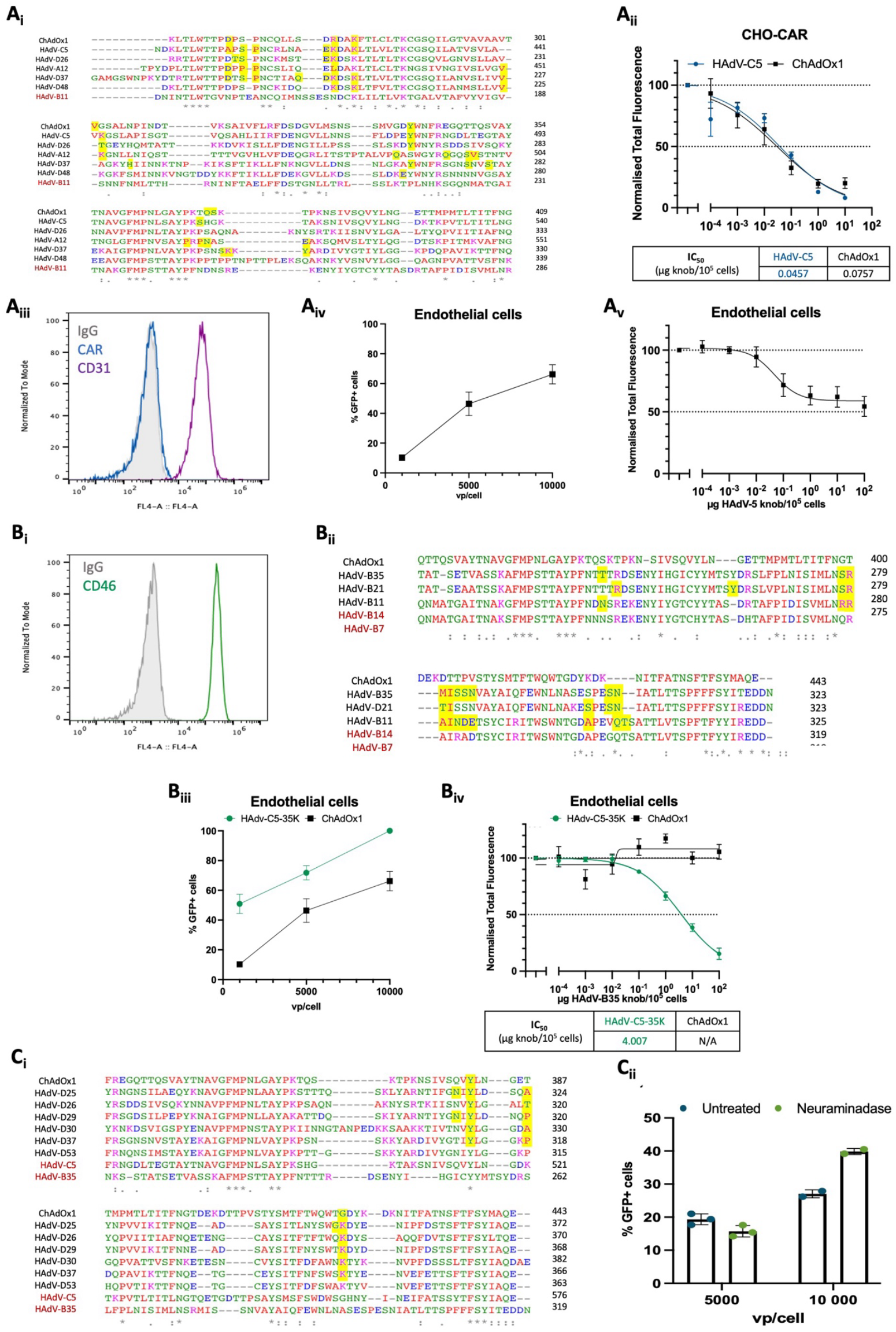
ChAdOx1 infects CAR-negative endothelial cells. (Ai) Clustal Omega sequence alignment of the ChAdOx1 fibre knob protein with other known CAR-binding HAdV fibre knob sequences[6]. Known CAR-binding residues are highlighted in yellow and demonstrate that ChAdOx1 shares many interacting residues with other CAR-binding adenoviruses. HAdV-B11 does not bind CAR. (Aii) Competitive inhibition studies in CHO-CAR cells, which express CAR but no other known primary HAdV receptors. Increasing concentrations of recombinant HAdV-C5 knob protein inhibited infection of the CHO-CAR cells by HAdV-C5-GFP and ChAdOx1-GFP, suggesting that ChAdOX1 is using CAR as a primary cell entry receptor. (Aiii) Immortalised human vascular endothelial cells are positive for CD31, a marker of endothelial cells, and negative for CAR. (Aiv) Transduction assays with ChAdOx1-GFP demonstrated infection of endothelial cells, suggesting the use of an unidentified cell entry receptor in the absence of CAR. (Av) Transduction of endothelial cells with ChAdOx1 was partially inhibited by HAdV-C5 fibre knob protein, suggesting that the predominant mechanism by which ChAdOx1 is infecting endothelial cells is independent of HAdV-C5. (Bi) Endothelial cells are positive for CD46, a primary cell entry receptor for some species B and D acenoviruses. (Bii) ChAdOx1 fibre knob protein sequences was aligned with other known CD46-binding HAdVs using clustal omega (Madeira *et al*. 2022). ChAdOx1 shared no CD46 binding residues, suggesting ChAdOx1 is not using CD46 as a cell entry receptor. HAdV-B14/B7 do not bind CD46. (Biii) Competitive inhibition studies in CHO-BC1 cells, which express CD46 but no other known primary HAdV receptors. Transduction of CHO-BC1 cells with HAdV-C5-35K-GFP (psuedotyped HAdV-C5 with HAdv-B35 knob protein) was inhibited by pre-incubation of the cells with recombinant HAdV-B35 fibre knob protein (IC_50_ 0.375 μg/10 cells). (Biv) HAdV-C5-35K and ChAdOx1 transduced 100% and 46% of endothelial cells respectively. (Bv) Transduction of endothelial cells with HAdV-C5-35K-GFP could be inhibited by pre-incubation with recombinant HAdV-B35 fibre knob protein (IC_50_ 1.043 μg/10 cells) whereas ChAdOx1 transduction was not inhibited, suggesting ChAdOx1 does not use CD46 to enter cells. (Ci) Using clustal omega sequence alignment (Madeira *et al*. 2022), the sequence of the ChAdOx1 fibre knob protein was compared with other known sialic acid-binding species D HAdVs. Sialic acid interacting residues are shown in yellow boxes. ChAdOx1 shared a tyrosine residue (Y381) in the sialic acid binding regions identified in HAdV-D26,D29, D30,D37 and – D53. (Cii) Pre-treatment of A549 and SKOV3 cells with neuraminidase to cleave sialic acid decreased the transduction of HAdV-C5-26K cells at 2000 vp/cell, indicating the use of sialic acid for cell entry. (Ciii) Pre-treatment of endothelial cells with neuraminidase did not decrease ChAdOx1 transduction at 5000 and 10 000 vp/cell suggesting ChAdOx1 is using another cell entry mechanism.VP, virus particles.

CAR regulates mechanotransduction, including maintenance and modulation of vascular function in EC[11]. CAR expression was undetected on EC (Figure 1_iii_); however; transduction assays with ChAdOx1.GFP demonstrated efficient transduction, suggesting use of an unidentified cell entry receptor (Figure 1_iv_). Competitive inhibition using HAdV-C5 fiber knob demonstrated HAdV-C5 fiber knob protein does not effectively block ChAdOx1 infection (Figure 1_v_). EC downregulate CAR expression in the presence of inflammatory cytokines, suggesting a mechanism by which EC can decrease adenoviral infection at sites of inflammation[12]. Therefore, the ability of ChAdOx1 to infect EC in the absence of CAR suggests that direct infection of EC could contribute to the pathogenesis of VITT.

CD46 is a primary attachment receptor for species B adenoviruses. EC CD46 expression was high (98.6% positive cells; MFI 6534) (Figure 1b_i_); however, ChAdOx1 fiber knob does not share CD46 binding residues for species B adenoviruses[9,13,14] (Figure 1b_ii_). Competitive inhibition assays in EC using HAdV-C5 adenovirus pseudotyped with the fiber knob from HAdV-B35 (HAdV-C5-35K), and ChAdOx1 demonstrated HAdV-B35K inhibits EC transduction of HAdV-C5-35K (IC_50_ 4.007μg/10^5^ cells) but not ChAdOx1 (Figure 1b_iii_;1b_iv_), suggesting ChAdOx1 does not engage CD46 to transduce EC (Figure 1b_v_).

Sialic acids (SA) are primary receptors for HAdV-Ds[15]. ChAdOx1 fiber knob sequences were aligned with clustal omega (Figure 1c_i_). Y381 (ChAdOx1), conserved across all sequences, is a key interacting residue for SA binding across HAdV-Ds, in addition T386 was shared with HAdV-D26. However, a key lysine residue of the SA binding pocket shared by HAdV-Ds is glycine (G420) in ChAdOx1. To investigate ChAdOx1 ability to engage SA, neuraminidase was used to remove cellular SA[15]. In EC, neuraminidase treatment did not decrease ChAdOx1 transduction at either viral concentration tested, suggesting ChAdOx1 does not use SA to enter cells (Figure 1c_ii_).

The ability of ChAdOx1 to transduce EC suggests a dual tropism where ChAdOx1 can transduce cells in the absence of CAR. Dual tropisms have been demonstrated in other HAdVs, such as the use of both CAR and sialic acid by HAdV-Ds[15,16]. Although this mechanism is still unclear, ChAdOx1 may utilise integrins to infect CAR-negative EC. Following attachment of HAdVs to cellular receptors, αvβ3 and αvβ5 integrins facilitate viral uptake through interaction of the penton base RGD motif. αvβ3 integrin is necessary for efficient infection of epithelial cells by HAdV-D26[17], suggesting ChAdOx1 infection may similarly be integrin-mediated. Alternatively, ChAdOx1 could engage heparan sulfate proteoglycans (HSPGs); HAdV-C2, HAdV-C5, HAdV-B3 and HAdV-B35 all demonstrate low affinity interactions with HSPGs, functioning as alternative receptors for infection[18,19]. Furthermore, other direct mechanisms of interaction exist between the ChAdOx1 capsid and the cell surface, such as the hexon-CD46 interaction demonstrated in HAdV-D56[20]. Investigation into the mechanism utilised by ChAdOx1 to transduce EC is required, and whether this interaction would be sufficient for productive infection of EC *in vivo*.

### ChAdOx1 does not induce endothelial cell activation

During endothelial dysfunction, EC become activated, releasing pro-coagulants that contribute to thrombosis generation. EC activation has been demonstrated in VITT patients and addition of VITT patient serum to EC increased P-selectin expression and platelet adhesion, suggesting EC activation contributes to VITT pathogenesis[21]. Furthermore, viral infection induces endothelial cell activation[22]. The ability of ChAdOx1 to induce EC activation was quantified by measurement of pro-thrombotic factors in the serum of transduced EC (Figure 2). Transduction with ChAdOx1, HAdV-C5 and HAdV-C5-B35K did not significantly increase EC production of Factor IX, interleukin-6 (IL-6), IL-8, plasminogen activator inhibitor-1 (PAI-1), P-selectin, glycoprotein ligand (PSGL), sCD40L and tissue factor (TF) compared to uninfected cells. Furthermore, expression of SARS-CoV-2 spike protein did not increase endothelial cell activation, further supporting that the activated endothelial cell phenotype in VITT patients is an indirect effect of ChAdOx1.

**Figure 2.**
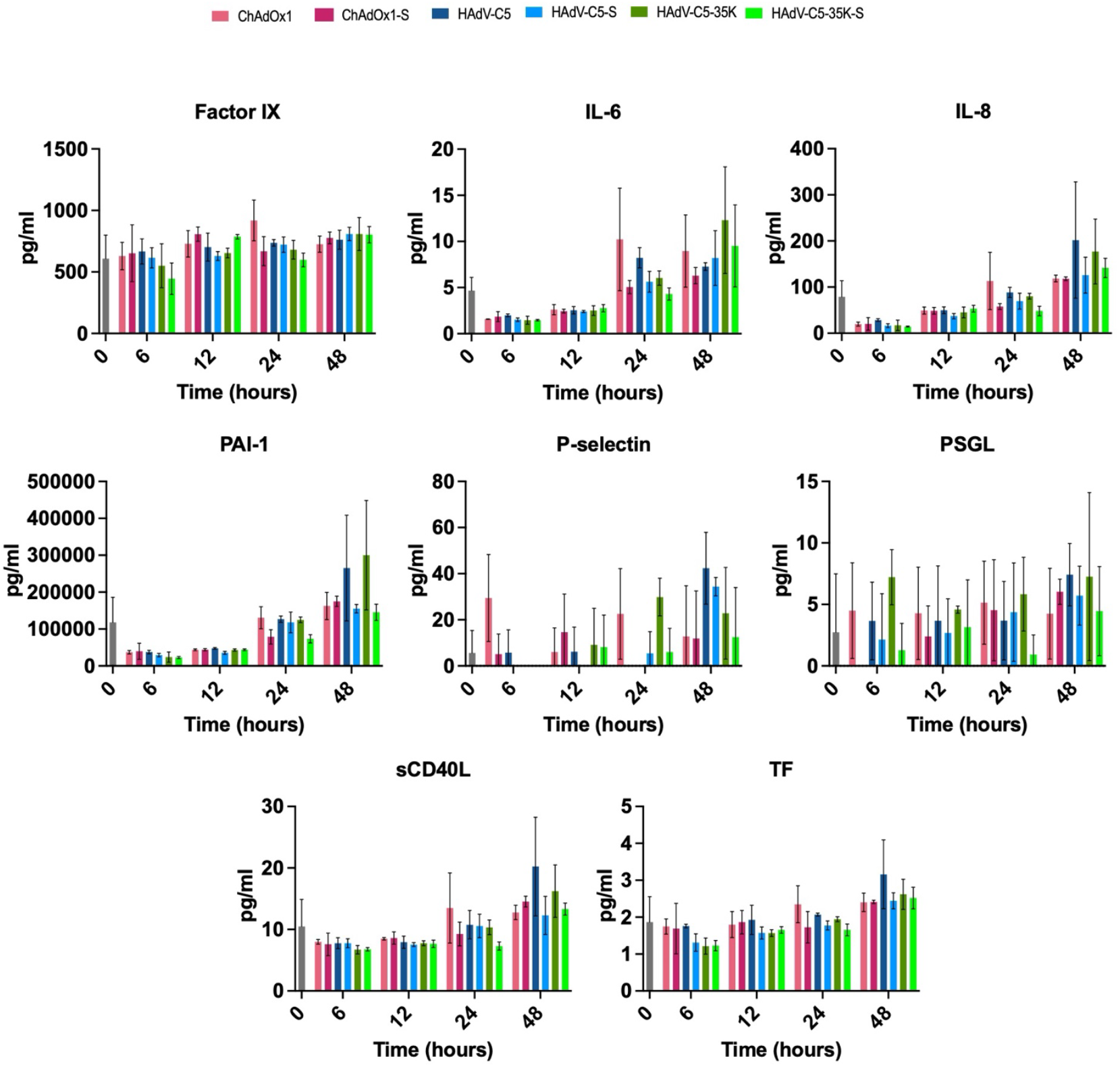
ChAdOx1 infection does not induce EC activation. Activated endothelial cells release pro-coagulants that contribute to thrombosis formation. ECs were infected with ChAdOx1, ChAdOx1-spike(S), HAdV-C5, HAdV-C5-S, HAdVC5-35K and HAdVC5-35K-S and the supernatants from infected cells were collected and analysed for EC thrombosis markers using the LEGENDplex human thrombosis panel. Infection with all viruses did not significantly increase EC production of any of these markers compared to uninfected cells (0h;grey bar) suggesting that endothelial cell activation in VITT is ChAdOx1- and spike protein – independent.

### Adenovirus hexon proteins bind PF4

Using SPR, we previously demonstrated that PF4 binds with nanomolar affinity to pure ChAdOx1 (KD 616 nM) and Vaxzevria preps (KD 514nm), as well as purified HAdV-D26, the platform of the Johnson & Johnson SARS-CoV2 vaccine, Jcovden [4]. These similar affinities confirm the interaction between the virus and PF4, rather than cell-line derived impurities in the vaccine[23]. A recent publication cast doubt on these findings, indicating that low pH acid washing during regeneration cycles caused increased PF4 binding, suggestive of low pH compromising adenoviral capsid stability, exposing negatively charged DNA, and promoting electrostatic binding of positively charged PF4 protein to viral DNA[24]. To address this concern, we generated purified preps of adenoviral hexon protein and performed SPR studies to assess direct PF4: hexon protein interactions in the absence of viral DNA. SPR analysis using immobilised purified hexon protein from ChAdOx1 demonstrated a greater affinity (KD 296.2 nM) for PF4 than that observed for both purified ChAdOx1 and Vaxzevria (Figure 3a), and higher affinity than HAdV-C5 hexon (KD 766.7 nM). However, the orientation of purified hexon protein exposes regions which would not otherwise be surface exposed on intact virions, suggesting such interactions could be non-specific charge-dependent interactions. Collectively, these data support direct electrostatic PF4 interactions with the ChAdOx1 viral capsid, rather than a direct infection and activation of EC by ChAdOx1 as the initiating event for VITT pathogenesis.

**Figure 3.**
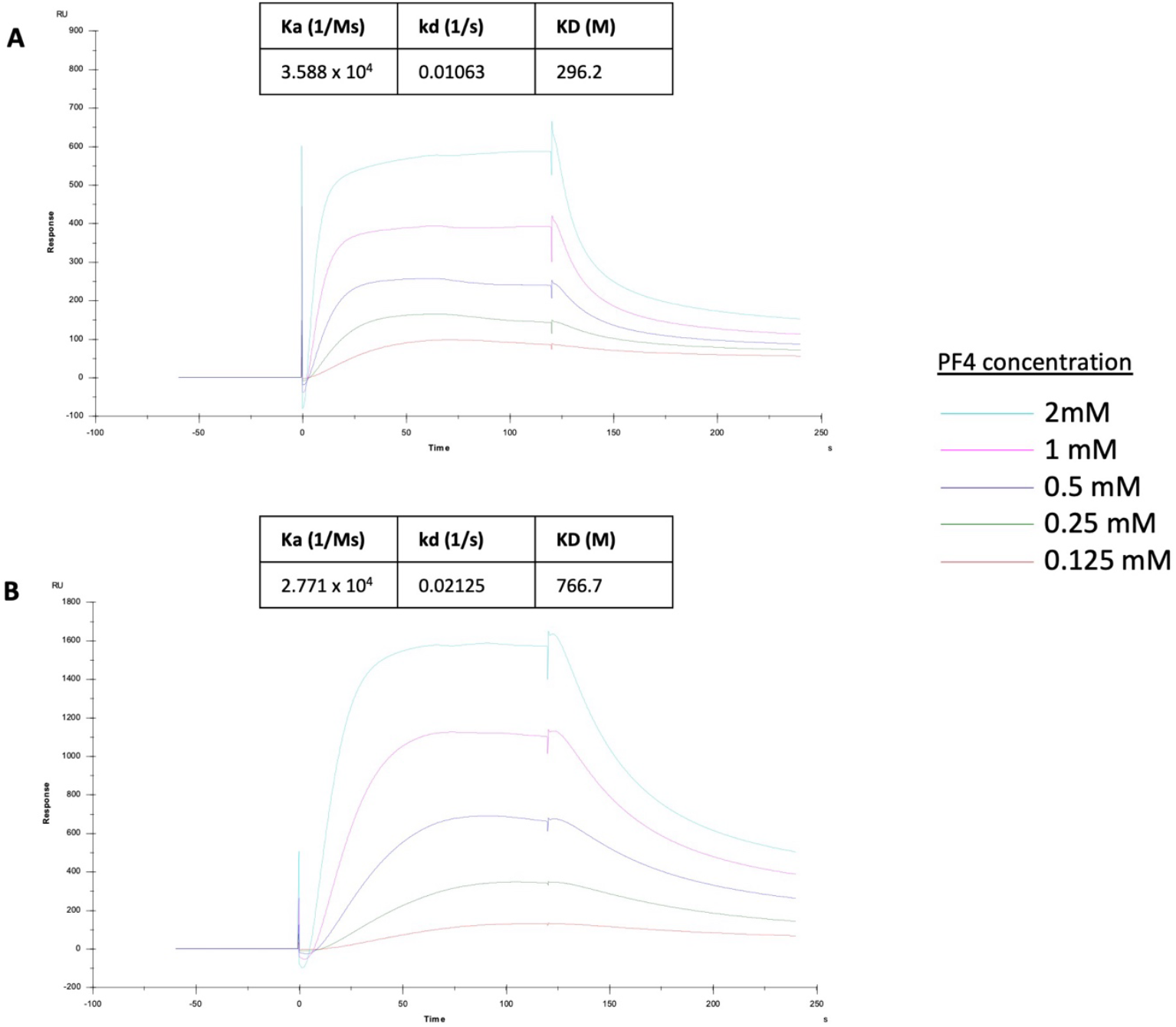
Adenovirus hexon proteins bind PF4. SPR analysis using PF4 at increasing concentrations over immobilised (A) ChAdOx1 and (B) HAdV-C5 hexon proteins demonstrates that ChAdOx1 hexon has a greater affinity for PF4 than HAdV-C5, and that previously reported for pure ChAdOx1 and Vaxzervia. This further supports that electrostatic interactions between PF4 and the ChAdOx1 viral capsid contribute to VITT pathogenesis. However, the orientation of purified hexon protein may expose regions which would not be surface exposed on intact virions, suggesting such interactions could be non-specific charge-dependent interactions.

## Conclusions

The ChAdOx1 nCoV-19 vaccine is safe and efficacious in protecting against symptomatic COVID-19 infection. Despite ultrarare occurrence of VITT, adenoviral vector-based vaccines remain vital worldwide. This study demonstrates that ChAdOx1 transduces CAR-negative EC, but this is unlikely to contribute to the pathogenies of VITT. Furthermore, the occurrence of VITT symptoms 5-24 days post-vaccination is consistent with anti-PF4 memory B cell responses. Furthermore we determine clear binding kinetics for PF4 binding purified ChAdOx1 hexon protein, confirming the interactions detected between PF4 and adenoviral vaccines vectors are not the result of low pH capsid destabilisation resulting in exposure of viral DNA. To develop safer adenoviral vaccines for the future, the focus should be on modification of the vector to prevent PF4 binding, or identification of adenoviruses which do not bind PF4.

## Authorship contributions

CL designed and conducted experiments, analysed data, and wrote the manuscript; LF designed and conducted experiments, analysed data, and reviewed the manuscript; EAS generated recombinant fiber knob proteins, and reviewed and edited the manuscript; RMM conducted experiments and reviewed and edited the manuscript; ALP designed experiments, acquired funding, supervised the study and wrote the manuscript.

